# Methadone blockade of I_K1_ promotes both long QT and Brugada-like arrhythmias: Mechanistic insights from computational modeling

**DOI:** 10.1101/2025.03.01.641012

**Authors:** Zhaoyang Zhang, J.T. Green, Mark C. Haigney, Kalyanam Shivkumar, Alan Garfinkel, Zhilin Qu

## Abstract

**Background:** Methadone is widely used for chronic pain relief and in the maintenance therapy of opioid use disorder, however, it also increases the risk of ventricular arrhythmias and sudden cardiac death. Methadone blocks several ionic currents with different half-maximal inhibitory concentrations (IC_50_), including the rapid component of the delayed rectifier potassium current (I_Kr_), the inward rectifier potassium current (I_K1_), the L-type calcium current (I_Ca,L_), and the late component of the sodium current (I_NaL_). Despite the well-known proarrhythmic effect of I_Kr_ blockade, the effects of blocking other ionic currents on arrhythmogenesis remain less well understood.

**Methods:** Computer simulations were used to explore the proarrhythmic effects of methadone by investigating how its blocking effects on ionic currents act alone or together in arrhythmogenesis.

**Results:** The major findings are: 1) blocking I_K1_ potentiates QT prolongation-related arrhythmogenesis by enhancing a tissue-scale dynamical instability for the spontaneous genesis of ectopic excitations. Blocking I_K1_ and I_Kr_ together results in a synergistic effect, greatly increasing the arrhythmia propensity, much larger than that of blocking either one alone; 2) blocking I_K1_ in combination with lowering I_Ca,L_ potentiates phase-2 reentry caused by spike-and-dome action potential morphology, an arrhythmia mechanism of Brugada syndrome. Blocking I_Kr_ exhibits little effect for this mechanism of arrhythmias; and 3) hypoxia, often comorbid in methadone populations, can potentiate QT prolongation-related arrhythmias at high sympathetic activity and phase-2 reentry at low or basal sympathetic activity, mainly via its effect on I_Ca,L._

**Conclusions:** These simulation results provide mechanistic insights into the genesis of QT prolongation-related Torsades de Pointes and Brugada-like ECG related arrhythmias caused by methadone use.

## Introduction

Methadone is a drug widely used for chronic pain relief and in the maintenance treatment of opioid use disorder, however, it is also associated with increased risk of ventricular arrhythmias and sudden cardiac death ^1-7^. Methadone users may exhibit abnormal ECG signs, including QT prolongation, Brugada-like ECG patterns, elevated U waves, etc.

Experimental studies have shown that methadone blocks the rapid component of the delayed rectifier potassium (K^+^) current (I_Kr_) ^8-13^, encoded by the human ether-a-go-go-related gene (hERG). Blocking I_Kr_ causes drug-induced long QT syndrome (LQTS) and Torsades de pointes (TdP), which agrees with methadone’s proarrhythmic profile of QT prolongation and TdP promotion.

However, biophysical studies have shown a wide range of half-maximal inhibitory concentrations (IC_50_) of methadone on I_Kr_, from 3 to 10 μM ^8-13^. In an analysis of clinical trial data ^14^, <5% of patients manifested peak methadone concentrations exceeding 3 μM. Due to extensive protein binding, free methadone concentrations may well be <1 μM. If the IC_50_ is much higher than 1 μM, the I_Kr_ blocking effect may not be as large as anticipated. A recent study by Klein et al ^13^ showed that methadone also blocks the inward rectifier K^+^ current (I_K1_) with an IC_50_ of 1.5 μM, suggesting that its blocking effect on I_K1_ may play a key role in arrhythmogenesis. In addition to blocking I_Kr_ and I_K1_, experimental studies have shown that methadone also blocks the L-type Ca^2+^ current (I_Ca,L_) and the late component of the sodium (Na^+^) current (I_NaL_) at high concentrations (5.5 μM and 8.5 μM, respectively) ^12, 15^, which agrees with the observations that methadone shortens action potential (AP) duration (APD) at high doses ^13^.

A recent study by Solhjoo et al ^16^ showed that methadone users have lower oxygen levels during sleep, which may cause hypoxia-induced electrophysiological changes to affect arrhythmogenesis. Despite the well understood role of blocking hERG in prolonging QT interval and promoting TdP, how the multiple ion channel blocking effects of methadone integrate together to impact arrhythmogenesis remains to be elucidated ^17^. Moreover, it is unclear why most methadone-related deaths occur during sleep ^1, 16, 18^.

In this study, we use computer simulations to investigate the mechanisms of arrhythmogenesis caused by methadone. Our major findings are: 1) Blocking I_K1_ potentiates APD- or QT prolongation-related arrhythmogenesis by enhancing the tissue-scale dynamical instability for spontaneous PVCs. Blocking I_K1_ and I_Kr_ together results in a synergistic effect, greatly increasing the arrhythmia propensity, much larger than that of blocking either one alone; 2) Blocking I_K1_ in combination with lowering I_Ca,L_ potentiates phase-2 reentry (P2R) caused by spike-and-dome AP morphology. Blocking I_Kr_ exhibits little effect for this mechanism of arrhythmias; and 3) Hypoxia, often comorbid in methadone populations, for example with underlying sleep apnea, can potentiate APD prolongation-related arrhythmias at high sympathetic activity and P2R at low or basal sympathetic activity, mainly via its effect on I_Ca,L_. We discuss implications of the mechanistic insights from the computer simulations to arrhythmogenesis caused by methadone.

## Methods

### PVCs in tissue

Early afterdepolarizations (EADs) in isolated cardiac myocytes are often cited as the cause of tissue-level PVCs, which then propagate through ventricular tissue and become triggers of arrhythmias. However, our computer simulations showed that while cellular-level EADs can cause tissue-level PVCs under the right conditions ^19, 20^, repolarization gradient (RG)-induced PVCs arising from tissue-scale dynamical instabilities might be the major mechanism of PVCs, which is supported by experimental observations ^21-24^. These PVCs, dubbed “R-from-T” ^25, 26^, can degenerate spontaneously into reentrant arrhythmias under the right tissue conditions. Therefore, to assess the proarrhythmic effects of methadone, we looked for PVC induction in tissue as our marker for arrhythmogenesis.

### 1D Cables

Based on our previous work, we found that a one-dimensional (1D) cable is a sufficient test for the genesis of these tissue-level PVCs and allows fast enough computer simulations. In our simulations, the 1D cable was composed of 200 cells (3 cm), the first half endocardium and the second half epicardium. We used four AP models, i.e., the 1994 Luo and Rudy (LRd) guinea pig model ^27^, the 2004 ten Tusscher at al (TP04) human model ^28^, the O’Hara et al (ORd) human model ^29^, and the modified ORd model by Tomek et al. (ToRORd) ^30^. The endocardial versions of the human models were used. For simplicity, we altered the slow component of the delayed rectifier K^+^ channel (I_Ks_) to model the endo-epi heterogeneity. We also added I_to_ to the epicardium.

### Populations of Models

To take into account inter-individual variabilities, we generated 1D cable model populations using random parameters and preset filters. The LRd, TP04, and ORd models were used for the model populations. (The ToRORd model behaves very similarly to the ORd model.) We multiply the maximum conductance of the major ionic currents with a factor drawn randomly from a preset interval. We carried out 1D cable simulations with the randomly selected parameter sets and measured APD at both the endocardium and epicardium. We first dropped the cases that exhibited PVCs and the cases that gave rise to too short an APD in the epicardium (due to a too strong I_to_). After these filters, we obtained a model population with a wide raw APD distribution. We then assigned a Gaussian distribution (the QTc distribution of healthy human is close to a Gaussian distribution ^31^) within this raw APD distribution for the endocardium as the target APD distribution, and then dropped parameter sets randomly to achieve the targeted Gaussian APD distribution. The retained parameter sets then form the normal control model population.

### Modeling channel block

The blocking effect of methadone on an ionic current was modeled by multiplying the current by a Hill function, i.e., for an ionic current I_m_, the formulation is:

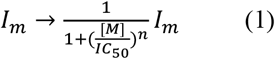

where [M] is the methadone concentration, IC_50_ is the half-maximal inhibitory concentration, and n is the Hill coefficient. I_m_ represents I_Kr_, I_K1_, I_Ca,L_, or I_NaL_. For I_Kr_, we used n=0.8 and IC_50_=2.9 μM (Klein et al ^13^ and Tran et al ^12^). For I_K1_, we used n=0.7 and IC_50_=1.5 μM (Klein et al ^13^). For I_Ca,L_, we used n=1.1 and IC_50_=5.5 μM (Tran et al ^12^). For I_NaL_, we used n=0.9 and IC_50_=8.5 μM (Tran et al ^12^).

## Results

### Effects of blocking I_Kr_ or I_K1_ on PVC genesis

To relate the effects of blocking I_Kr_ or I_K1_ on APD to their effects on PVC genesis, we first explored the effects of blocking I_Kr_ or I_K1_ on APD for the four AP models simulated in this study (Fig.1). In all four models, reducing I_K1_ by 50% results in roughly a 5% increase in APD but reducing I_Kr_ by 50% causes a larger APD increase, as expected. However, their responses to blocking I_Kr_ are very different, with the ORd and ToRORd models exhibiting a much larger change in APD than did the other two models.

**Figure 1.**
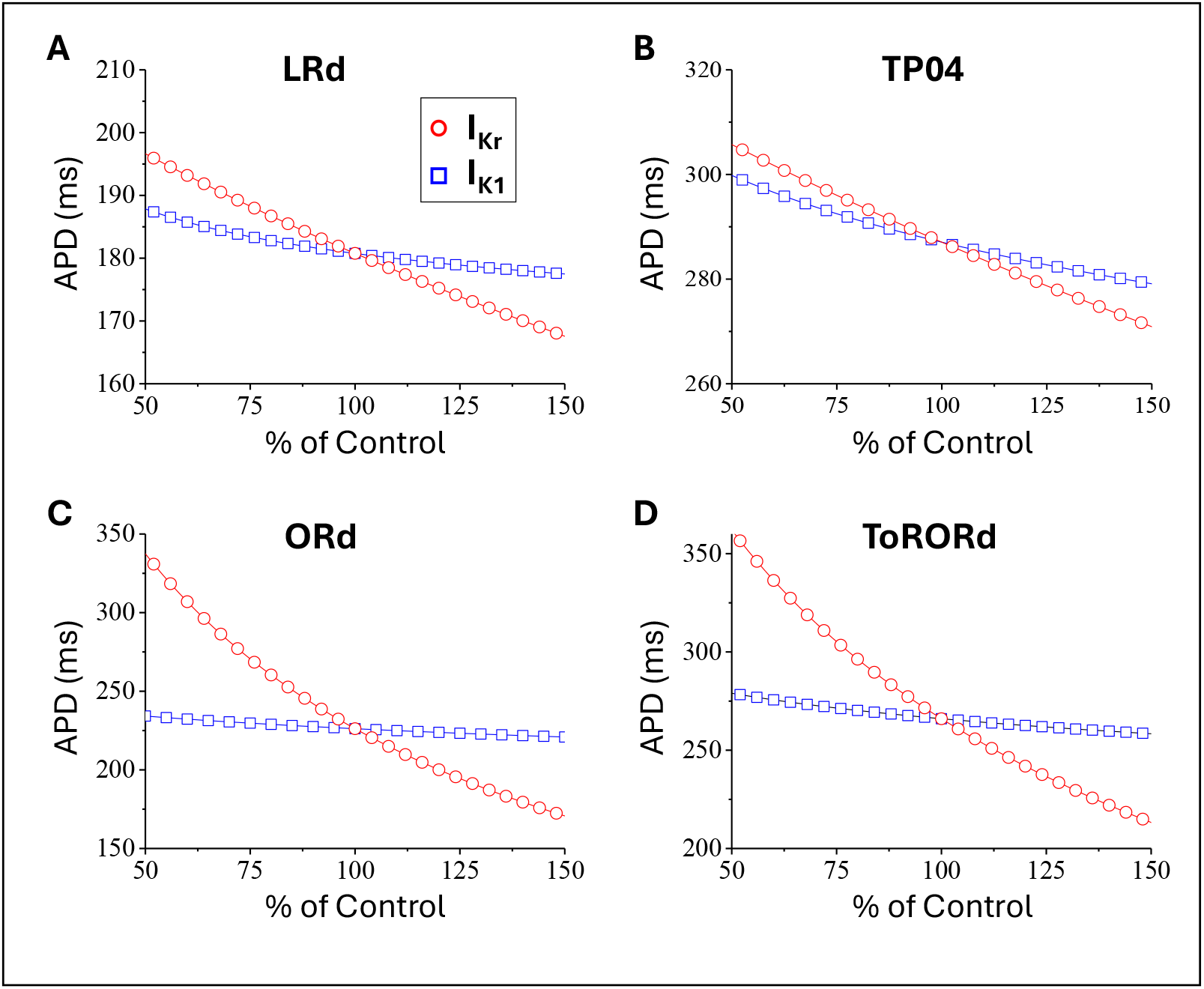
Effects of blocking I_Kr_ and I_K1_ on APD prolongation. Showing are APD versus percentage of the control value of G_Kr_ (circle) or G_K1_ (square) for the LRd model (A), the TP04 model (B), the ORd model (C), and the ToRORd model (D).

To investigate the effects of blocking I_Kr_ or I_K1_ on PVC genesis, we performed simulations of 1D cables, scanning for what combinations of the maximum conductance of I_Kr_ or I_K1_ and I_Ca,L_ produced PVCs. To model repolarization heterogeneity, we reduced G_Ks_ in the endocardial region and added I_to_ in the epicardial region. We first used cables based on the LRd model to show PVC genesis in more detail (Fig.2) and then cables based on TP04, ORd, and ToRORd models (Fig.S1) to verify and compare with the LRd model results.

#### EAD- and Repolarization Gradient-mediated PVCs

For the LRd model, we first excluded I_to_ in the epicardial layer to avoid I_to_-mediated PVCs (Fig.2A). In this case, PVCs are promoted by enhancing I_Ca,L_ and blocking I_K1_ or I_Kr_ (light blue regions). In other words, for a fixed G_K1_ or G_Kr_, PVCs occur once G_Ca,L_ is above a certain threshold, or for a high enough G_Ca,L_, PVCs can occur by reducing G_K1_ or G_Kr_. These PVCs are mediated by early afterdepolarizations (EADs) in the long APD region or by the Repolarization Gradient (RG) between endocardium and epicardium (see an example of RG-mediated PVC in Fig.2D), as shown in previous studies ^19, 20^. Although blocking I_K1_ increases APD less than does blocking I_Kr_ (Fig.1A), it causes a faster reduction of the threshold I_Ca,L_ for PVC genesis than blocking I_Kr_. For example, a 50% reduction of I_Kr_ from its control value reduces the I_Ca,L_ threshold from 4.5 to 3.9 fold (red arrow in right panel in Fig.2A) of its control value, while a 50% reduction in I_K1_ reduces the threshold from 4.5 to 3 fold (red arrow in left panel in Fig.2A). *This indicates that although I*_*K1*_ *only exhibits a small effect on APD and EADs at the cellular scale* ^*32*^, *it plays a critical role at the tissue scale to promote PVCs. This is because both EAD- and RG-mediated PVCs are tissue-scale phenomena driven by tissue-scale dynamical instabilities* ^19, 20^, *and I*_*K1*_ *plays a critical role in promoting these dynamical instabilities in tissue*.

#### Phase-2 Reentry-mediated PVCs

We then added I_to_ to the epicardial layer and observed qualitative difference in PVC genesis (Fig.2B). Adding I_to_ created a spike-and-dome AP morphology in the epicardial layer (Fig.2C) but had only a small effect on RG-mediated PVCs, i.e., the light blue regions at high G_Ca,L_ in Figs. 2 A and B are similar. However, a new PVC region emerged at low G_Ca,L_ (magenta regions and insets in Fig.2B). The mechanism of this type of PVCs is via phase-2 reentry (P2R) caused by the I_to_-induced spike-and-dome AP morphology ^33-35^. As shown in Fig.2E, the AP in the epicardial layer became very short (a spike), facilitating the formation of a PVC. The PVC region of this mechanism reduces as G_K1_ increases until it completely disappears. In other words, blocking I_K1_ promotes P2R. In general, P2R is promoted by decreasing inward currents or increasing outward currents. It is non-intuitive that blocking I_K1_, an outward current, promotes P2R. This is because similar to the RG-mediated PVCs, the PVCs caused by P2R are also a result of a tissue-scale instability ^36^ that depends on I_K1_ strength. Note that the P2R region is insensitive to I_Kr_ changes (the width of the PVC region only changes slightly with G_Kr_), agreeing with that shown in our previous studies that P2R is not sensitive to changes of G_Kr_ and G_Ks_^35, 37^.

**Figure 2.**
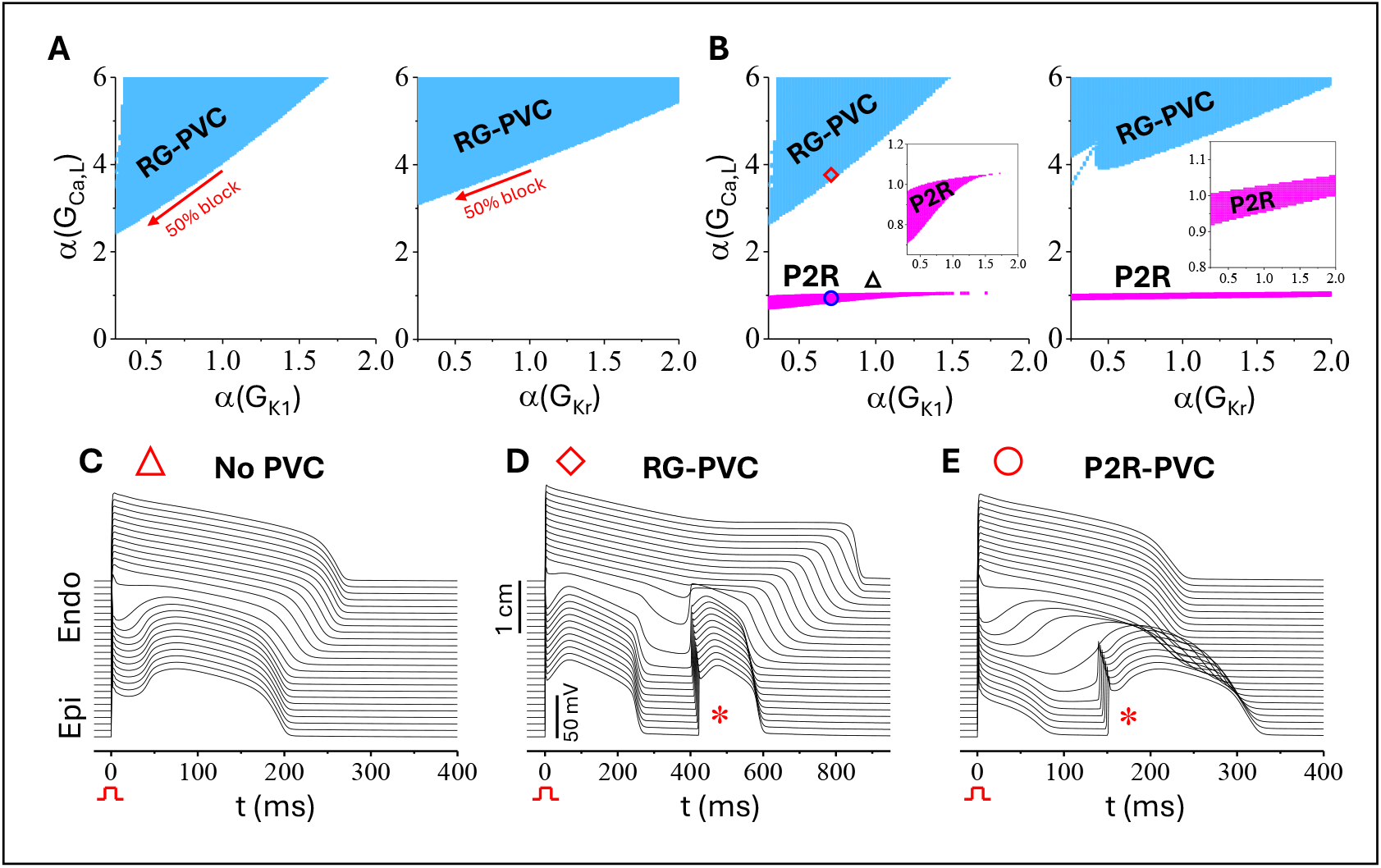
Effects of I_K1_ and I_Kr_ on PVC genesis in a 1D cable with the LRd model. **A**. RG- mediated PVC regions (light blue) in the G_K1_ and G_Ca,L_ parameter plane (left) and the G_Kr_ and G_Ca,L_ parameter plane (right) when no I_to_ is present in the epicardial region (G_to_=0). G_Ks_ is 0.3 folds of control in the endocardium and 2 folds in the epicardium. Red arrows mark the changes in I_Ca,L_ threshold for a 50% block of I_Kr_ from control. **B**. Same as A but when I_to_ is present in the epicardial region (G_to_=1.75 mS/cm^2^). The RG-mediated PVC regions are colored light blue and the P2R- medtaed PVC regions are colored magenta. The three symbols mark the parameter locations for the examples shown in C-E. Insets are expanded views of the P2R-mediated PVC regions. **C**. Time-space-voltage plot of a case with no PVC (triangle, G_K1_ is 1 fold and G_Ca,L_ 1.1 folds of their control values). **D**. Time-space-voltage plot of a RG-mediated PVC caused by reducing I_K1_ and increasing I_Ca,L_ (diamond, G_K1_ is 0.7 folds and G_Ca,L_ 3.7 folds of control). * marks the spontaneous PVC. **E**. Time-space-voltage plot of P2R-mediated PVC caused by reducing both I_K1_ and I_Ca,L_ (circle, G_K1_ is 0.7 folds and G_Ca,L_ 0.9 folds of control). * marks the spontaneous PVC. Note that the stimulated AP in the epicardial layer is an abbreviated spike due to lowering G_Ca,L_, which allows the spike-and-dome AP in the middle layer to initiate a PVC in epicardial layer.

As shown in our previous simulation studies of 2D tissue ^19, 35, 37^ or 3D heart ^25^ models, the RG-mediated or P2R-mediated PVCs can maintain as multi-beat focal excitations or degenerate into reentrant arrhythmias, depending on the tissue conditions. Therefore, in this study, we used the 1D cable model as our test platform, and the genesis of spontaneous PVCs as a marker for arrhythmogenesis.

We compared the results of the LRd model to those of the other three AP models (see SI Fig.S1). The results are similar despite some quantitative differences.

In summary, blocking I_Kr_ and/or I_K1_ combined with increasing I_Ca,L_ conductance promotes RG-mediated PVCs, and blocking I_K1_ combined with decreasing I_Ca,L_ conductance promotes P2R- mediated PVCs. Although blocking I_K1_ only exhibits a small effect on prolonging APD and promoting EADs in single cells, it exhibits a much larger effect on promoting PVCs in tissue, a tissue-scale phenomenon that cannot be revealed by single-cell studies.

### Arrhythmogenesis promoted by methadone

In general, we can infer, from the results shown in Figs.1, 2 and S1, that among the proarrhythmic effects of methadone, blocking I_Kr_ and I_K1_ prolongs APD and promotes arrhythmias under high I_Ca,L_ via RG-mediated PVCs and that blocking I_K1_ promotes arrhythmias at low I_Ca,L_ via P2R-mediated PVCs. However, methadone also blocks other currents and causes hypoxia during sleep; how these effects of methadone integrate together to affect arrhythmogenesis is unclear.

Secondly, in the general population, there is substantial inter-individual variability: individual parameter sets will differ from one person to another. This inter-individual variability results in individuals having different responses to a specific drug, including different responses to different drug doses. The results shown in Figs.1 and 2 are based on a single set of control parameters for each model, which is not representative of the diversity of the general population. To take into account inter-individual variability, we generated model populations using randomly selected parameter sets and APD filters (see Methods). This randomization-plus-filtering resulted in Gaussian-like distributions of APD from the endocardium of the normal control populations for the LRd model (Fig.3A), the TP04 model (Fig.S2A), and the ORd model (Fig.S3A). To differentiate the contributions of methadone’s blocking effects on different ionic currents, we also used different combinations of its blocking effects. For example, we might include only its I_K1_ or I_Kr_ blocking effect alone or a combination of the two, etc.

#### Effects of Methadone on APD

We first used the model populations to assess the effects of methadone on APD under normal control conditions. Fig.3B shows average APDs and standard deviations for different methadone concentrations and different combinations of methadone blocking effects, marked by different colors, for the LRd model. In general, the responses of the average APD of the model populations (see also SI Figs. S2B and S3B) to methadone agree with the results shown in Fig.1: 1) blocking I_K1_ alone (yellow) only lengthens APD slightly; 2) blocking I_Kr_ alone (red) has a bigger effect on prolonging APD than does blocking I_K1_; 3) blocking I_K1_ and I_Kr_ together (blue) results in stronger effects on lengthening APD than did blocking I_Kr_ alone; and 4) the APD lengthening effects due to blocking I_K1_ and I_Kr_ are attenuated by blocking I_Ca,L_ and I_NaL_ (green and purple).

#### Arrhythmogenesis in high sympathetic conditions

We then used the model populations to simulate the arrhythmogenic effects of methadone in the setting of high sympathetic activity, modeled by increased G_Ca,L_. We chose to modify only G_Ca,L_ because, although β-adrenergic stimulation increases the amplitudes of both I_Ca,L_ and I_Ks_, the amplitude of I_Ca,L_ increases much faster than that of I_Ks_^38^, and thus in the early phase of sympathetic stimulation, the main effect is the increase of I_Ca,L_ strength. Fig.3C shows the percentages of individuals in the population exhibiting PVCs for the LRd model. When methadone only blocks I_Kr_ (red bars), the PVC risk, although it increases with methadone dose, is very low. If methadone blocks I_K1_ only (yellow bars), the risk increases with dose to a higher percentage than by blocking I_Kr_ alone. Interestingly, when methadone blocks both I_Kr_ and I_K1_ together (blue bars), the risk becomes much higher, indicating that blocking I_Kr_ and I_K1_ synergistically promotes arrhythmias. The TP04 model (Fig.S2C) behaves similarly to the LRd model. The ORd model (Fig.S3C) exhibits some differences from the other two models. In the ORd model, blocking I_K1_ exhibits almost no effects while blocking I_Kr_ exhibits a large effect. But again, blocking both I_Kr_ and I_K1_ together exhibits a much larger effect, demonstrating that the synergistic effects of blocking both I_Kr_ and I_K1_ still occur as observed in the other two models. For all three models, as anticipated, blocking I_Ca,L_ reduces PVCs (green bars), which is substantiated at high doses of methadone. Due to its high IC_50_, the effect of I_NaL_ on PVC genesis is small (purple bars) with a larger effect occurring in the ORd model.

To correlate APD to PVC risk under different conditions of methadone blocking effects, we plotted the percentage of the population exhibiting PVCs versus averaged baseline APD for different conditions of methadone blocking effects (Fig.3D). Specifically, we plotted the percentage exhibiting PVCs (under high sympathetic activity) against the averaged APD shown in Fig.3B (under control). Blocking I_Kr_ alone lengthens APD but does little in promoting PVCs (red). Blocking I_K1_ alone increases APD slightly but has a large effect on promoting PVCs (yellow, steep curve). Blocking both I_K1_ and I_Kr_ together, the curve is less steep than for blocking I_K1_ alone but extends to longer APDs and higher percentages due to the synergy effect. The TP04 model behaves similarly to the LRd model (Fig.S2D) while the ORd model behaves differently when blocking I_Kr_ or I_K1_ alone (Fig.S3D). However, all three models demonstrated that blocking I_Kr_ and I_K1_ together greatly enhanced the arrhythmia risk.

#### Arrhythmogenesis in low sympathetic conditions

Finally, we simulated the effects of methadone under low sympathetic activity (control I_Ca,L_) for the LRd model (Fig.4) and the TP04 model (SI Fig.S4). We simulated two scenarios: with or without hypoxia. Since we simulated low sympathetic activity, we used the effects of hypoxia on ionic currents under basal condition used by Gaur et al ^39^, in which I_Ca,L_ was reduced. In the absence of hypoxia (Fig.4A), when methadone blocks I_K1_ and/or I_Kr_ but does not block I_Ca,L_ and I_NaL_, it exhibits almost no effects on PVC genesis (color bars do not show up in graph). When methadone also blocks I_Ca,L_ and I_NaL_ (green and purple bars), the PVC risk increases with methadone concentration quickly due to its I_Ca,L_ blocking effect. Blocking I_NaL_ only enhances the risk slightly. In the presence of hypoxia (Fig.4B), blocking I_K1_ increases the arrhythmia risk (yellow) but blocking I_Kr_ with or without blocking I_K1_ does not affect the risk of arrhythmia (red and blue bars). If methadone also blocks I_Ca,L_ and I_NaL_ (green and purple bars), the PVC risk decreases with methadone concentration due to too high an I_Ca,L_ blockade at high methadone concentrations.

#### *Role of I*_*ca*_,_*L*_

Since I_Ca,L_ is the current affected by sympathetic stimulation, hypoxia, and methadone, it plays a key role in methadone-induced arrhythmias. To give broader view of the effects of I_Ca,L_ combined with I_K1_ and I_Kr_ blocking effects, we recorded the percentage of PVCs versus I_Ca,L_ strength for a fixed methadone concentration (3 μM) with methadone blocking I_Kr_ (red boxes) or I_K1_ (yellow boxes) alone or both together (blue boxes) for the LRd model (Fig.5). For reference, we also plot the percentage of PVCs in the absence of methadone. In all cases, the PVC risk is low in the intermediate I_Ca,L_ strength range. For high I_Ca,L_, PVCs are caused by APD prolongation, and the risk increases with the strength of I_Ca,L_. For low I_Ca,L_, PVCs are caused by spike-and-dome AP morphology via P2R, which occurs in a narrow range of I_Ca,L_ strength. Blocking I_K1_ greatly increases the risk of PVCs, but blocking I_Kr_ exhibits little effects.

## Discussion

In this study, we used computational simulation to investigate the multiple ion channel blocking effects of methadone on arrhythmogenesis, beyond its role as a hERG blocker. Our major findings are:

1. Blocking I_K1_ does not prolong APD significantly but can substantially increase the occurrence of RG-mediated PVCs at high sympathetic activity via tissue-scale instabilities. Blocking I_K1_ works synergistically with blocking I_Kr_ to greatly increase the susceptibility to arrhythmias. Blocking I_Ca,L_ (and I_NaL_) at high methadone concentrations can attenuate the risk of arrhythmias, as anticipated.
2. Blocking I_K1_ promotes P2R at low or basal sympathetic activity, which is potentiated by the I_Ca,L_ blocking action of methadone. Blocking I_Kr_ has little effect on this mechanism of arrhythmias.
3. Hypoxia may promote RG-mediated PVCs at high sympathetic activity by enhancing I_Ca,L_ and I_NaL_ and also promote P2R at basal or low sympathetic activity by suppressing I_Ca,L_.

The insights from the computer simulations reveal the important roles of I_K1_ in both RG-mediated and P2R-mediated PVCs and the mechanisms of methadone-induced arrhythmogenesis, as discussed below.

## I_K1_ blockade and its synergy with I_Kr_ blockade in APD prolongation and arrhythmogenesis

It is well understood that blocking I_Kr_ lengthens APD and promotes EADs, which are associated with QT prolongation and arrhythmogenesis in the inherited LQTS type 2 and most cases of drug-induced LQTS. Blocking I_K1_ gives rise to a small increase in APD (Fig.1) and thus its contribution to QT prolongation is small.

But on the other hand, our present and previous studies ^19, 20^ showed that although blocking I_K1_ plays only a small role in APD prolongation (Fig.1) and in EAD genesis ^32^, it plays a much greater role in promoting PVCs in tissue (Figs.2 and 3). We showed that the RG-mediated (or phase-2 EAD-mediated) PVCs originate from a tissue-scale dynamical instability caused by RG and enhanced I_Ca,L_^19, 20, 36^, and I_K1_ potentiates this instability without affecting the APD or the RG per se. *Therefore, the main effect of I*_*K1*_ *on PVC genesis is at the tissue level by promoting dynamical instability in tissue, while the effect of I*_*Kr*_ *is mainly at the single-cell level by prolonging APD and promoting EADs*.

### Effect on DADs

Because I_K1_ plays a key role in stabilizing the resting potential, reducing I_K1_ increases the amplitude of DADs and promotes DAD-mediated triggered activity creating arrhythmogenesis ^40, 41^.

### Phase-3 EADs

For the same reason, I_K1_ is thought to be important for the genesis of phase- 3 EADs ^17^, which have been observed in Purkinje fibers ^42-44^ and in mouse and rat ventricular myocytes ^45, 46^. However, our survey of the literature showed that no phase-3 EADs had been observed in isolated ventricular myocytes of large animals and humans ^32^. Computer simulations showed that to generate phase-3 EADs in ventricular myocytes of large animal or human models, drastic changes in ionic current conductance are needed ^47^, and these large changes in parameters may well be unphysiological. Therefore, whether blocking I_K1_ promotes TdP via promoting phase- 3 EADs or not remains unclear.

More intuitively, reducing I_K1_ makes the resting potential or phase-3 repolarization of the AP less stable, allowing spontaneous occurrence of depolarizations (thus PVCs) in the RG region. Note that phase-3 EADs may be rarely observed in isolated ventricular myocytes but can be observed in tissue ^22^ due to electronic coupling and the tissue-scale genesis of PVCs. For example, in Fig.2D, one can record a phase-3 EAD in the middle of the cable (the RG region) while there are no EADs in the long and short APD regions at all.

### Synergy between I_K1_and I_Kr_ blockade

Another important finding is that blocking I_K1_ works in synergy with blocking I_Kr_ to promote PVCs. In the cases shown in Figs.3, S2 and S3, when only I_Kr_ was blocked, increasing the methadone concentration from 0 to 5 μM increased the PVC risk from 0 to 0.18% in the LRd model, to 0% in the TP04 model, and to 4.1% in the ORd model. When only I_K1_ was blocked, the PVC risk increased from 0% to 9.36%, 3.58%, and 0%, respectively. However, when both were blocked, the PVC risk increased from 0% to12.56%, 23.54%, and 20.06%, respectively. Therefore, blocking both I_Kr_ and I_K1_ causes an increase in PVC risk that is disproportionate to the sum of blocking them alone, indicating a strong synergistic effect.

**Figure 3.**
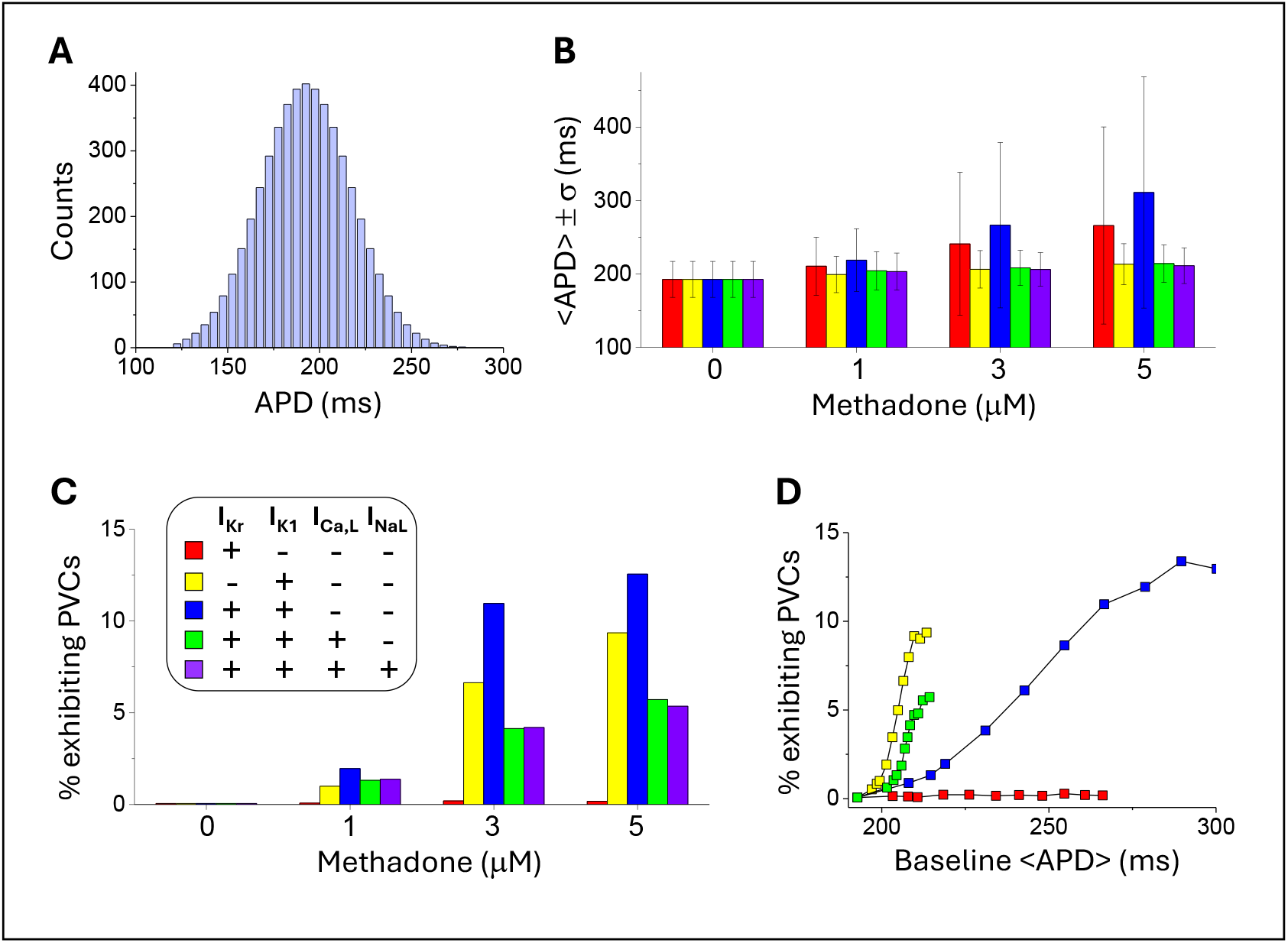
Effects of methadone on APD and arrhythmia risk in the LRd model. **A**. APD histogram of the model population under the control condition. **B**. Bar graph showing averaged APD (<APD>) and standard deviation (σ) versus methadone concentration. **C**. Bar graph showing percentage of individuals exhibiting PVCs versus methadone concentration under 5 different conditions of methadone blocking effects (see labels in the inset). The I_Ca,L_ is strength is 5 folds of control for the LRd model. **D**. Dependence of percentage of individuals exhibiting PVCs on baseline <APD> under the first 4 conditions of methadone effects. The baseline <APD> was obtained at control I_Ca,L_ as in B and the PVCs are obtained with the same I_Ca,L_ as in C.

The synergy can be understood as follows. Blocking I_Kr_ prolongs APD and enhances RG while blocking I_K1_ does not prolong APD or enhance RG but destabilizes the system. When the two work together, they work cooperatively or synergistically, amplifying the effect on PVC genesis. This synergistic effect may explain the experimental ^48^ and clinical ^49^ observations that when I_Kr_ and I_K1_ are reduced simultaneously, arrhythmia events increase disproportionately compared to reducing either one alone. Specifically, in the study by Hornyik et al ^48^, they showed that the arrhythmia rate was <5% for transgenic LQT2 or LQT2-5 rabbits under control but when treated with an I_K1_ blocker, the rate increased to >50%. However, there were little effects in the wild type or LQT5 rabbits with the same treatment. In the clinical study by Mazzanti et al ^49^, they showed that in Andersen-Tawil Syndrome Type 1 patients (where Kir2.1 mutation reduces I_K1_), treatment with the I_Kr_ blocker amiodarone (a class III antiarrhythmic drug) caused a >40-fold increase in arrhythmia risk compared to treatments with β-blockers or class 1c antiarrhythmic drugs.

## I_K1_ and I_Ca,L_ blockade in P2R-mediated arrhythmogenesis

P2R is a term used to describe a PVC or reentry caused by the spike-and-dome morphology induced by I_to_^33, 34^ or by altered Na^+^ channel kinetics ^37^. It has been proposed as one mechanism of arrhythmogenesis in Brugada syndrome (BrS). In general, BrS and P2R are promoted by reducing inward currents and/or increasing outward currents, in contrast to LQTS. For example, our simulations showed that P2R is promoted by increasing I_to_ (Figs. 2 and S1) or by reducing I_Ca,L_ (Fig.5). A non-intuitive observation from our simulation is that blocking I_K1_, an outward current, can greatly increase the risk of PVCs (Figs. 4, 5, and S4). Blocking I_K1_ also promotes P2R caused by altered Na^+^ channel kinetics ^37^. Here, the role of I_K1_ is the same as its role in promoting RG-mediated PVCs, i.e., by promoting tissue-scale dynamical instabilities ^36^ or by making the resting potential and phase-3 repolarization less stable.

**Figure 4.**
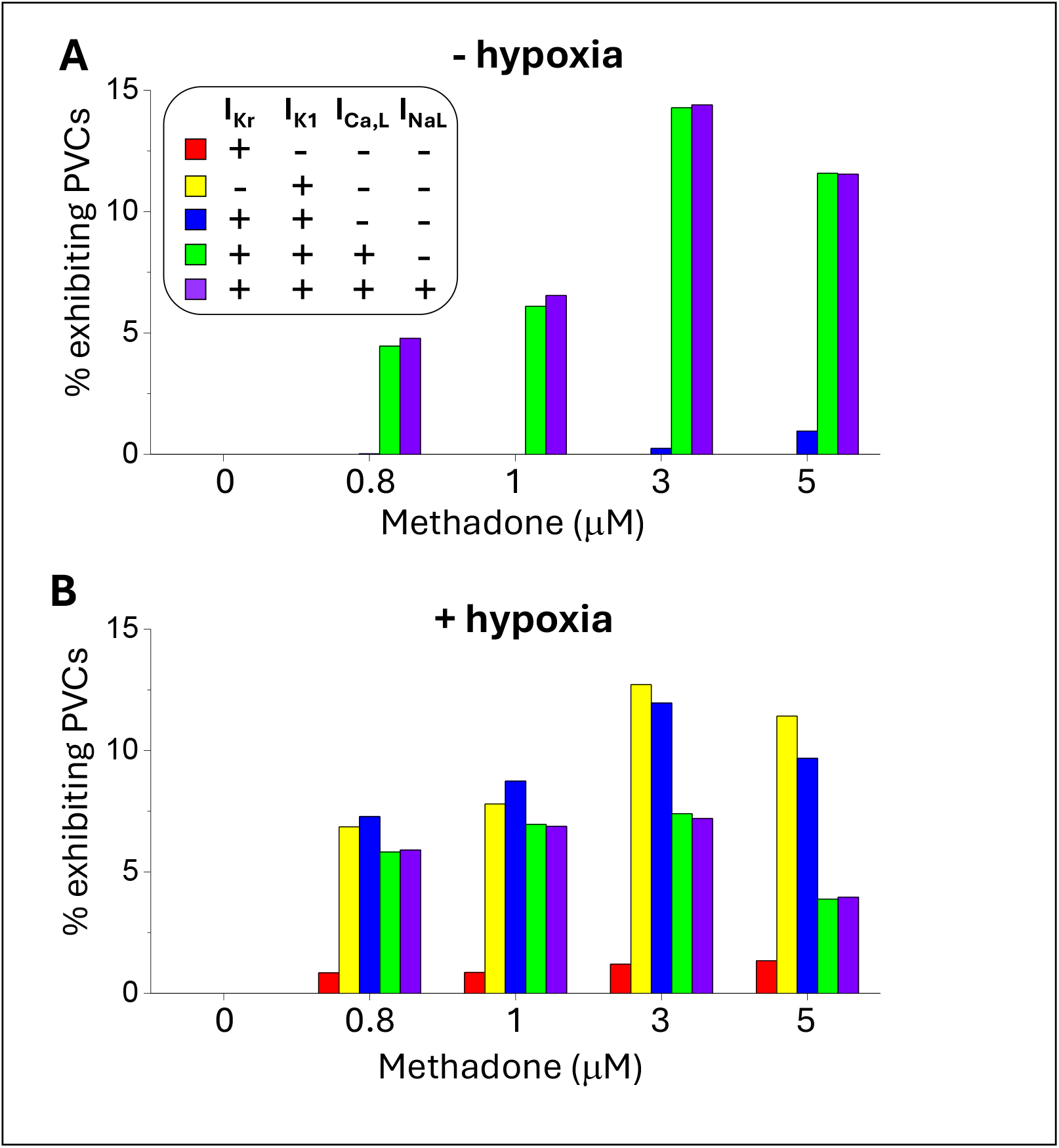
Effects of methadone on PVC genesis (P2R) at low sympathetic activity (low I_Ca,L_ strength) in the LRd model. **A**. Percentage of individuals in the population exhibiting PVCs versus methadone concentration under 5 different conditions of methadone blocking effects (see labels in the inset) for the control G_Ca,L_ without hypoxia. **B**. Same as A but for the control G_Ca,L_ with hypoxia. Effects of hypoxia on ionic current conductance are modeled as follows (From Gaur et al with zero isoproterenol ^39^): *I*_*Na*_ → 0.9*I*_*Na*_, *I*_*NaL*_ → 1.5*I*_*NaL*_, *I*_*Ca,L*_ → 0.75*I*_*Ca,L*_, and *I*_*Ks*_ → 0.73*I*_*Ks*_.

**Figure 5.**
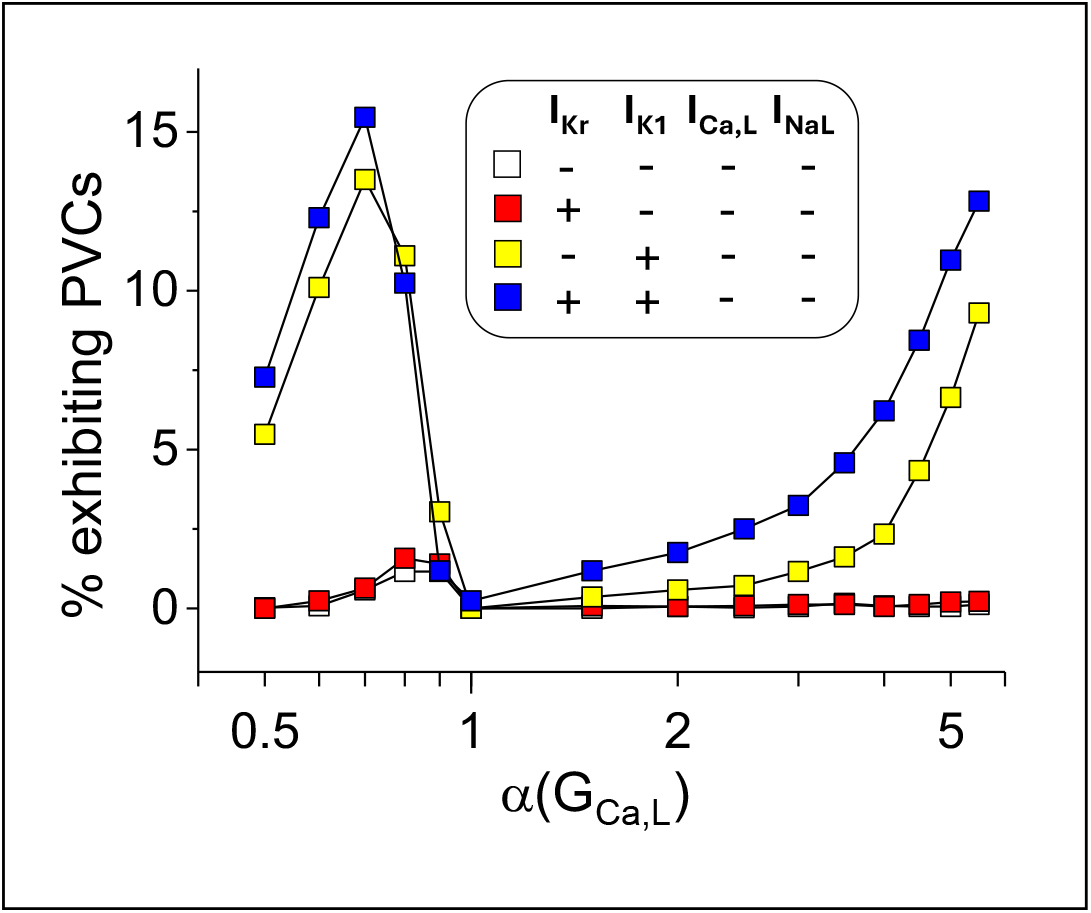
Percentage of PVCs versus I_Ca,L_ strength. Shown are percentage of individuals exhibiting PVCs versus G_Ca,L_ for the LRd model under no methadone action and three different conditions of methadone effects as labeled in the inset. The methadone concentration is 3 μM. The open boxes almost completely overlap with the red boxes.

## Roles of hypoxia in QT prolongation and P2R mediated arrhythmogenesis

Based on the mathematical formulation by Gaur et al ^39^, hypoxia alters I_Na_, I_NaL_, I_Ca,L_ and I_Ks_ and their responses to isoproterenol. For example, in the mathematical formulation by Gaur et al ^39^, *I*_*Ca,L*_ → *Isofactor* × *I*_*Ca,L*_ with 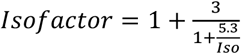 for the normal condition and 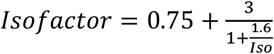 for hypoxic condition. For low isoproterenol (e.g., for Iso=0, Isofactor=1 for the normal condition and 0.75 for hypoxia), I_Ca,L_ is lower under hypoxia than under the normal condition. For intermediate or high isoproterenol (e.g., for Iso=1.6, Isofactor=1.69 for the normal condition and 2.25 for hypoxia), I_Ca,L_ is higher under hypoxia than under the normal condition. Therefore, at low sympathetic activity or basal condition, hypoxia reduces I_Ca,L_, but at normal or high sympathetic activity, it increases I_Ca,L_. The consequence is that hypoxia promotes arrhythmias at both low and high sympathetic activity via promoting P2R-mediated and RG-mediated PVCs, respectively.

## Arrhythmogenetic effects of methadone

QT prolongation has been shown for methadone use ^13, 14, 50, 51^ with high risk of TdP ^7^. Like many other LQTS or TdP-genic drugs, methadone also blocks I_Kr_ and thus causes QT prolongation and TdP as widely anticipated. However, biophysical studies showed a wide range of IC_50_, from around 3 to 10 μM ^8, 10-13^ or even higher ^9^. For example, for a methadone concentration of 1 μM and an IC_50_ 3 μM, I_Kr_ is only blocked by ∼25%. If the IC_50_ becomes 10 μM, only ∼10% I_Kr_ is blocked. Therefore, the I_Kr_ blocking effect may not be sufficient to explain methadone’s proarrhythmic effect.

On the other hand, it has been shown that methadone blocks I_K1_ with an IC_50_ of 1.5 μM ^13^. Our simulations showed that blocking I_K1_ alone, although it only has a small effect on prolonging APD or the QT interval, can increase the risk of RG-mediated PVCs. More interestingly, blocking I_K1_ and I_Kr_ simultaneously results in a synergistic effect which increases the risk of arrhythmias many-fold more than blocking either one alone. In other words, moderately blocking I_Kr_ combined with blocking I_K1_ increases QT moderately, but the synergistic effect of blocking the two greatly increases the TdP risk, making methadone a drug causing a high incidence of TdP with moderate QT prolongation.

### Methadone and Brugada Syndrome-like arrhythmias

While the major focus of methadone effect is on QT prolongation and TdP, BrS-like ECGs have also been observed in methadone users ^5, 52-55^. Our simulations in this study showed that methadone’s I_K1_ blocking effect combined with its I_Ca,L_ blocking effect can promote P2R at low or basal sympathetic activity, agreeing with arrhythmogenesis in patients with BrS. Note that unlike QT prolongation, BrS-like ECGs are not persistent even in the same patient and thus cannot be easily or widely detected. Clinically, Na^+^ channel blockers are often used to unmask BrS. Furthermore, our simulation showed that blocking I_Kr_ prolongs QT but has little effect on either promoting or suppressing P2R, and thus QT prolongation can still be a dominant ECG marker, but the mechanism of arrhythmias can be P2R as for BrS. Therefore, the prevalence of BrS-related arrhythmias in methadone users might be highly underestimated.

### U-waves

Elevated U-waves are another ECG feature in methadone users ^5, 6^, which agrees with methadone’s I_K1_ blocking effect ^13, 56^. Although the exact mechanism of the U-wave is unclear, it is related to phase-3 repolarization and phase-4 depolarization, where I_K1_ becomes important. One of the links of U-waves to arrhythmias is via DAD-mediated arrhythmogenesis ^57, 58^ in which reducing I_K1_ lowers the threshold for a DAD to trigger an AP for arrhythmogenesis ^40, 59^. DADs are caused by spontaneous intracellular calcium release events, which can be triggered by high sympathetic activity ^60^.

### Hypokalemia

Besides its ion channel blocking effects, another arrhythmia risk factor for methadone use is hypokalemia ^6, 61-63^. One of the effects of hypokalemia is reduction of I_K1_ conductance, and lowering I_K1_ is a key parameter for arrhythmias in all three mechanisms, as discussed above. Therefore, in the case of methadone use, I_K1_ is already lowered by the drug action and hypokalemia makes it even worse.

### Effects of sleep

Finally, our simulation results may provide insights into the observation that most methadone deaths occur during sleep ^1, 16, 18^. It is known that cardiac arrhythmias for LQTS occur during physical or emotional stress except for LQT3 where the cardiac events occur mainly during sleep ^64, 65^. For catecholaminergic polymorphic ventricular tachycardia (CPVT), arrhythmias occur in the afternoon to evening due to higher physical activity ^66^, and for BrS, most arrhythmias occur in the nighttime due to low sympathetic activity ^67^. Our simulations show that blocking I_K1_ combined with lowering I_Ca,L_ caused by low sympathetic activity and methadone-related ion channel blockade and hypoxia during sleep promotes P2R for arrhythmogenesis. This agrees with the observations that BrS-like ECGs occur in methadone users and that methadone-related deaths mainly occur during sleep. Furthermore, serum K^+^ concentration is lower in the nighttime (bottoming around 9pm) ^68-70^, which can further increase the likelihood of arrhythmias during sleep.

## Limitations

Several limitations are worth mentioning. First, we simulated only 1D cable models for spontaneous genesis of PVCs, to allow for a large number of simulations, but arrhythmias are tissue-level phenomena in the three-dimensional heart. To model arrhythmias, two- or three-dimensional tissue models are needed. Furthermore, the shape of the geometry of the heterogeneous region in higher dimensions than 1D may also be nontrivial for arrhythmogenesis. However, as shown previously ^26, 35, 37^, once a spontaneous PVC occurs, it may exhibit as an ectopic beat or degenerate into reentrant arrhythmias, depending on properties of the heterogeneous region. Therefore, the propensity of spontaneous PVCs in the 1D cable is a good measure of propensity for arrhythmias in higher dimensional tissue. Second, we used one fixed IC_50_ for each current and combinations of inclusion and exclusion of currents for methadone action to assess arrhythmogenesis related to methadone use. In reality, the IC_50_ for methadone action may differ in different experimental conditions, in different species, as well as in different individuals. These differences may give rise to different conclusions for the contributions of different ionic currents caused by methadone. Third, we modeled sympathetic activity by altering I_Ca,L_ only. It is known that β-adrenergic stimulation causes strong increases of both I_Ca,L_ and I_Ks_. Since the maximum I_Ca,L_ strength elevates much faster than that of I_Ks_ in response to β-adrenergic stimulation ^38^, there is a time window in which I_Ca,L_ enhancement is the dominant effect. Fourth, although we simulated 4 AP models and obtained generally consistent results, there are still differences between models, such as the roles of I_Kr_ and I_K1_. Nevertheless, the computer simulation results in this study provide mechanistic insights into methadone-induced arrhythmogenesis and sudden cardiac death.

## Conclusions

Methadone blocks multiple ionic currents, which alone or in combination can promote arrhythmias via LQTS related arrhythmias (TdP) and BrS related arrhythmias (P2R). In particular, its I_K1_ blockade effect can work synergistically with its action of blocking other currents, such as I_Kr_ and I_Ca,L_, to greatly increase the propensity of arrhythmias and sudden cardiac death.

## Notes

### Competing Interest Statement

The authors have declared no competing interest.

